# RosettaSurf - a surface-centric computational design approach

**DOI:** 10.1101/2021.06.16.448645

**Authors:** Andreas Scheck, Stéphane Rosset, Michaël Defferrard, Andreas Loukas, Jaume Bonet, Pierre Vandergheynst, Bruno E Correia

## Abstract

Proteins are typically represented by discrete atomic coordinates providing an accessible framework to describe different conformations. However, in some fields proteins are more accurately represented as near-continuous surfaces, as these are imprinted with geometric (shape) and chemical (electrostatics) features of the underlying protein structure. Protein surfaces are dependent on their chemical composition and, ultimately determine protein function, acting as the interface that engages in interactions with other molecules. In the past, such representations were utilized to compare protein structures on global and local scales and have shed light on functional properties of proteins. Here we describe RosettaSurf, a surface-centric computational design protocol, that focuses on the molecular surface shape and electrostatic properties as means for protein engineering, offering a unique approach for the design of proteins and their functions. The RosettaSurf protocol combines the explicit optimization of molecular surface features with a global scoring function during the sequence design process, diverging from the typical design approaches that rely solely on an energy scoring function. With this computational approach, we attempt to address a fundamental problem in protein design related to the design of functional sites in proteins, even when structurally similar templates are absent in the characterized structural repertoire. Surface-centric design exploits the premise that molecular surfaces are, to a certain extent, independent of the underlying sequence and backbone configuration, meaning that different sequences in different proteins may present similar surfaces. We benchmarked RosettaSurf on various sequence recovery datasets and showcased its design capabilities by generating epitope mimics that were biochemically validated. Overall, our results indicate that the explicit optimization of surface features may lead to new routes for the design of functional proteins.

**Author Summary:** Finely orchestrated protein-protein interactions are at the heart of virtually all fundamental cellular processes. Altering these processes or encoding new functions in proteins has been a long-standing goal in computational protein design.

Protein design methods commonly rely on scoring functions that seek to identify amino acid sequences that optimize structural configurations of atoms while minimizing a variety of physics-based and statistical terms. The objectives of the large majority of computational design protocols have been focused on obtaining a predefined structural conformation. However, routinely introducing a functional aspect on designer proteins has been more challenging.

Our results suggest that the molecular surface features can be a useful optimization parameter to guide the design process towards functional surfaces that mimic known protein binding sites and interact with their intended targets. Specifically, we demonstrate that our design method can optimize experimental libraries through computational screening, creating a basis for highly specific protein binders, as well as design a potent immunogen that engages with site-specific antibodies. The ability to create proteins with novel functions will be transformative for biomedical applications, providing many opportunities for the design of novel immunogens, protein components for synthetic biology, and other protein-based biotechnologies.

## Introduction

Proteins are key components in living cells, performing many functions that commonly rely on physical interactions between them and other molecules. Steady progress in structural biology has unveiled the structures of a wide variety of proteins and their mode of interaction with other molecular partners. High-resolution protein structures are studied at the atomic level, focusing on the interactions between residues, backbone conformations, or the topology of secondary structure elements, among others (1). The molecular surface arising from the three-dimensional arrangement of the many atoms that compose a protein is, ultimately, determinant for protein function and is therefore crucial for biological processes (2).

While discrete atomic-level protein representations have been invaluable for our understanding of protein function, near-continuous surface-based representations offer the opportunity to study protein structures using a different subset of features (e.g. electrostatic potentials, geometry).

In 1971, Lee and Richards introduced the concept of solvent-accessible surfaces, which in practice are generated by rolling a probe approximating a solvent molecule over the protein atoms (3). The molecular surface, often denoted as solvent-excluded surface or Connolly surface (4–6), consists of the surface that can be directly contacted by the probe and the reentrant surface which smooths over gaps between the atoms that were not accessible to the probe sphere (5,7,8). Numerical representations of surfaces have also been developed, ranging from dot surfaces, to voxel representations and graphs (1,2,9–12). These representations allow the mapping of molecular and geometric properties onto the generated surface, including physicochemical properties (such as electrostatics and hydrophobicity), and geometrical features (e.g. protrusions or cavities) (1,7,13). Having these features of proteins encoded in their surfaces facilitates powerful computational analysis. Recently, the machine learning-based approach MaSIF (molecular surface interaction fingerprinting) showcased the extraction of surface features from molecular surfaces to study protein structure-function relationships (14,15).

Intuitively, the molecular surface forms the boundary of the protein and its surroundings, thus acting as the interface that engages in interactions with other molecules. The study of protein structures as near-continuous molecular surfaces is therefore important to understand structural and functional aspects of proteins, which may not be fully captured by a discrete atomic representation (13,16). A widely studied category of features of molecular surfaces is their chemistry, in particular electrostatic potentials, and their implications for protein function. Most notably, such representations have been used to study various types of protein-substrate interactions (17,18). However, many different aspects of proteins can be investigated with this approach, including the solvation energies of proteins in different media, redox potentials, or catalytic rates of enzymes (17). Other important metrics used to study molecular surfaces are their shape-derived properties, commonly focusing on shape complementarity (S_C_) or shape similarity (S_S_). Shape complementarity is frequent in the study of molecular recognition, e.g. protein-protein or protein-ligand interactions. In particular, protein-protein interactions (PPIs) have been extensively studied in terms of complementarity, showing that protein-protein interfaces are often highly complementary, both in shape and in charge (19,20). Moreover, shape complementarity has been incorporated into protein docking to guide the prediction of protein complexes (21–25) and to match binding pockets with potential ligands (26). On the other hand, shape similarity has been used to globally and locally compare biomolecules, aiding the functional annotation of proteins that show structural similarities but lack detectable sequence homology (11,27). Thus, shape similarity allows for the classification of local surface patches as well as the prediction of ligand binding sites by pocket shape similarity (1,11).

An important extension of such successful applications in analysis and prediction tasks using surface-based representations is that of protein design (7), where the objective is to guide the sequence search process to optimize specified surface features.

Here we present a new, surface-centric computational design strategy, termed RosettaSurf, that uses a description of the molecular surface shape and electrostatic properties as objective function for scoring optimization in protein design. Working at the surface-level of protein structures offers a unique approach for the design of proteins and their functions, as molecular surfaces are, to some extent, independent of the underlying sequence (11,14). This means that different sequences in different proteins may present similar surfaces (11,14) – a premise that drives our proposed methodological approach. In contrast, the large majority of computational design workflows entail a discrete atomistic description of the proteins and sequence design is typically performed on the atom arrangements and interactions; i.e. by sampling different amino acid side chains and adjusting atoms in the protein structure to optimize a given energy function. While these strategies have been successfully applied to design novel protein topologies (28– 30), the design of functional proteins based solely on computational calculations remains a challenge (31–38). The introduction of function into a designer protein often requires the transplantation of functional sites by means of protein grafting (39–44). A major limitation of this approach is the dependence on the identification of appropriate protein scaffolds (35). Therefore the ability to accurately design surface patches could represent an alternative route to functionalize computationally designed proteins.

RosettaSurf is a protocol implemented in the Rosetta software package (45) and we assessed its performance with several benchmark tests, demonstrating the protocol’s capabilities through the recovery of protein interfaces and its application to functional protein design.

## Results

In our benchmark studies we sought to compare the developed surface-centric design protocol, RosettaSurf, to state-of-the-art macromolecular design approaches implemented in Rosetta. The Rosetta energy function has been parametrized to evaluate and optimize the energy of many different aspects of molecular interactions (e.g. protein stability, protein-ligand, protein-protein and protein-nucleic acids, etc.); it contains both statistical and physics-based terms, being calibrated using a discrete atomic representation of the molecules. This type of optimization has been particularly successful for the design of novel sequences that fold into defined protein structures. However, to design proteins that display defined motifs which can perform biological functions has proven to be difficult. By focusing the design process on the areas where molecular interactions occur – the protein surfaces – novel approaches attempting to design function into proteins may represent new routes to address this problem.

Two types of metrics in surface comparison are considered: surface complementarity and surface similarity. We show how the surface similarity score captures features of individual residues and demonstrate the ability of the RosettaSurf protocol to recover amino acid sequences of native protein interfaces. Furthermore, we highlight the performance of RosettaSurf for the design of surface patches at protein interfaces.

### Single amino acid recovery

To evaluate the accuracy of our design protocol, we performed a benchmark on the recovery of single amino acids in native protein interfaces. The amino acid of interest is substituted by each of the 19 amino acid identities (excluding cysteine) and the surfaces of the substituted rotamers are compared to that of the native rotamer (Fig. 1A). The energy computed by Rosetta, Rosetta Energy Unit (REU), serves as a baseline for comparison to the surface-centric design strategy. We evaluated the recovery of the different amino acids for mutations made in the bound and unbound state of the protein complex, respectively (Fig. 1B). The recovery of an amino acid is deemed successful when the native amino acid has the best score among all 19 amino acids and is uniquely identifiable. In this benchmark we evaluated the performance of shape similarity, electrostatic similarity and both combined in a surface similarity score. The surface similarity measurement is highly accurate in identifying native amino acids in the bound state of the protein complexes, showing consistently higher sequence recovery rates than Rosetta energy (Fig. 1B). Incorporating the electrostatic similarity term into the surface score generally results in a boost in recovery rates over shape similarity alone, in particular for amino acids that have close structural doubles but differ in chemical properties, e.g. glutamine and glutamate, as well as, asparagine and aspartate (Fig. 1B).

**Figure 1.**
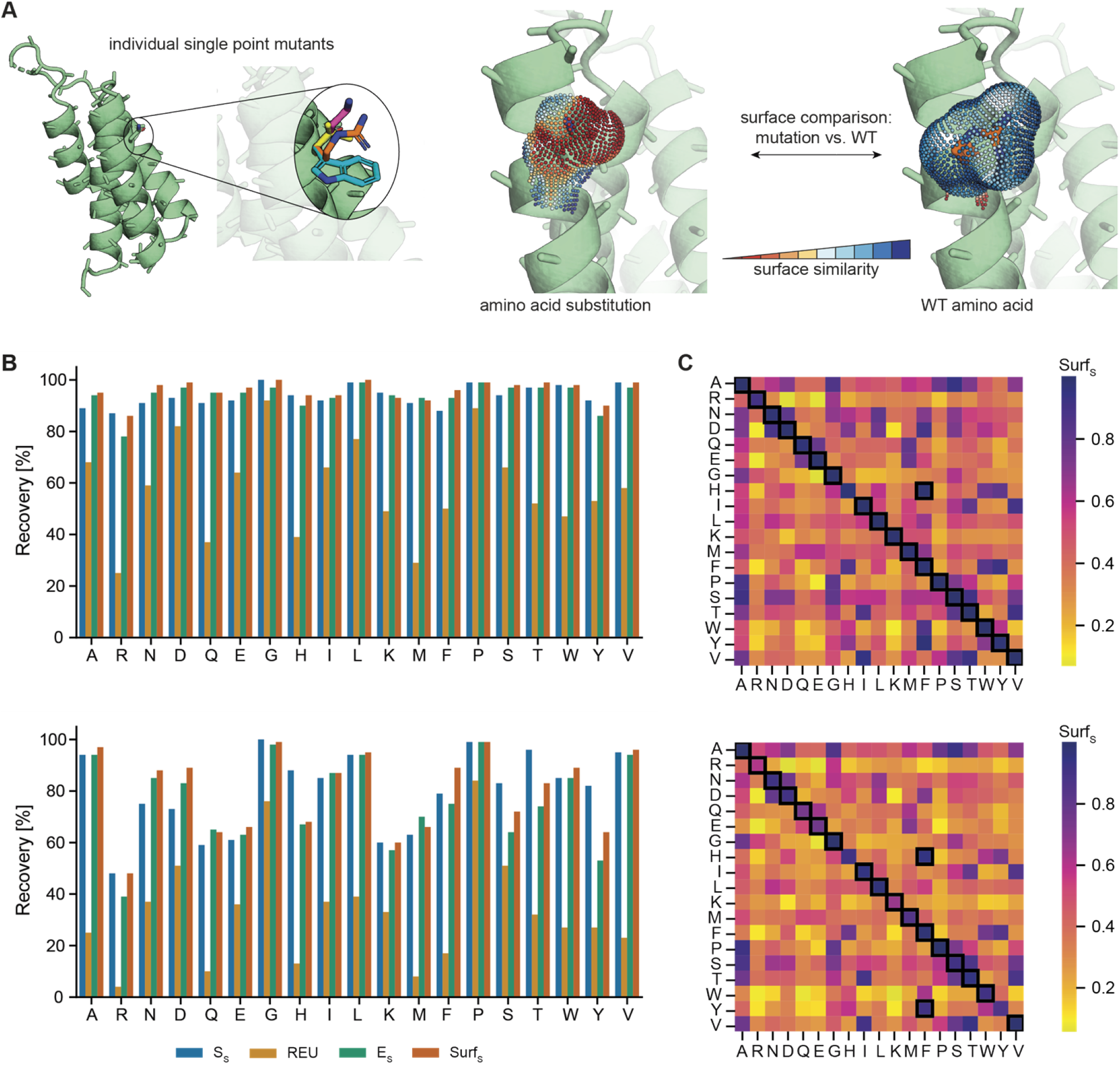
Single mutant discrimination using surface similarity score in protein-protein complexes. A) Surface similarity evaluation protocol for single amino acids. B) Recovery for all 19 considered amino acid types in bound (top) and unbound (bottom) complex states. C) Average surface similarity score when performing all-against-all amino acid comparison for bound (top) and unbound (bottom) complex states. The highest mean Surf_S_ score for every amino acid is highlighted.

The Rosetta energy function shows the best recovery rates for amino acids with unique features on their side-chains, i.e. glycine and proline. However, even for these cases, surface similarity outperforms the recovery by REU. A similar trend can be observed for the unbound complexes, although the general success of retrieving the native amino acid decreases for certain amino acids. This trend is most obvious for polar residues and large side chains that have access to a variety of different rotamer conformations as the lack of the binding partner allows a larger conformational space for different rotamers to be explored.

Moreover, the surface similarity scores retrieved from an all-against-all amino acid comparison demonstrate the high accuracy by which most amino acids can be identified (Fig. 1C). Here, we computed mean surface similarity values for all substitutions tested, i.e. for each native amino acid across the 100 protein complexes we computed the similarity of all other 19 amino acids to the native one on average. The results show that each of the 19 amino acids is generally most similar to itself, demonstrating that the method can accurately distinguish between the different amino acid types. Notable exceptions are residues that share similar geometrical features, e.g. phenylalanine and histidine or tyrosine.

Close inspection of the considered surface of a single amino acid demonstrates the local precision of the surface similarity score (Fig. S2). Small off-rotamer deviations are captured on the point cloud and are specific for the mismatching surface area. These results indicate that the surface similarity score is sensitive to small differences in the comparison of two surface point clouds. Overall, these results clearly show that the developed surface similarity score can capture local differences in surfaces providing the basis for the evaluation of differences between full surface patches.

### Protein interface sequence recovery

Having shown that the implemented surface similarity score is sufficiently accurate to recover individual amino acids, the following benchmark expands on recovering entire surface patches in protein interfaces. We evaluated the ability of our surface-based sequence design protocols (RosettaSurf and RosettaSurf-site) to recover natural protein interfaces as a gateway for the design of proteins endowed with biochemical function. By focusing on protein interfaces, the correct sequence does not solely depend on minimizing the Rosetta scoring function, but rather needs to represent the surface properties of the interface site. While such design scenario has limited applicability for the *de novo* design of protein-protein interactions, it is important for applications in the domain of immunogen design for vaccine development where the surface mimicry of known surfaces (neutralizing epitopes) is critical for the biological activity of this type of design (34,36–38).

In this benchmark we considered nine protein-protein complexes, grouped into three categories (low complementarity, high complementarity, antibody/antigen), and evaluated the performance by assessing sequence recovery (Fig. 2A). We compared three different design approaches:

**Figure 2.**
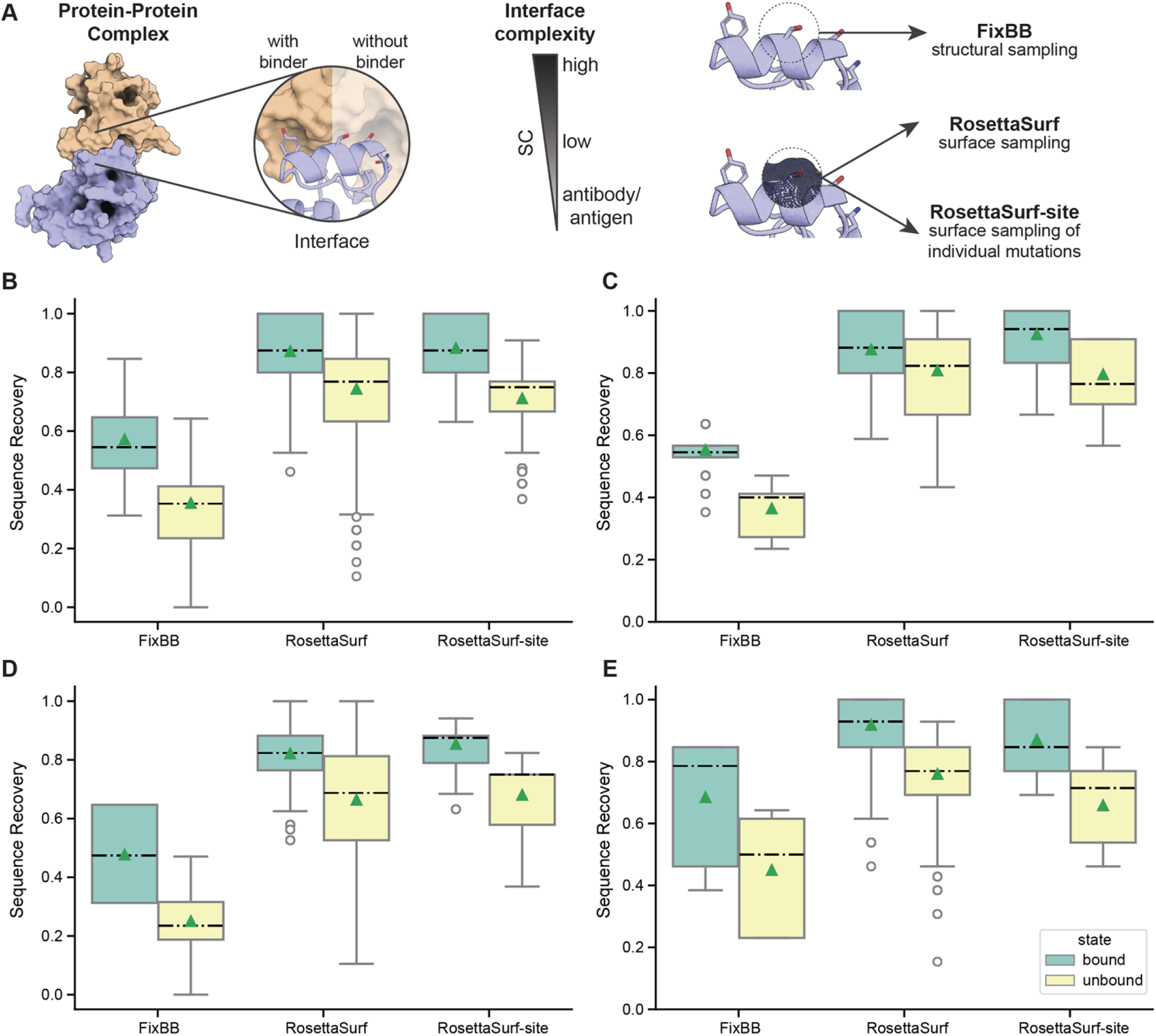
Sequence recovery of protein interfaces. A) Sequence recovery benchmark pipeline. Sequences in the interfaces of protein-protein complexes are evaluated in the presence and absence of the binding partner. The tested complexes were grouped into interfaces with low and high shape complementarity, and antibody-antigen complexes. Surface-centric design (RosettaSurf and RosettaSurf-site) is compared to a standard structural protein design protocol (FixBB). B) Interface sequence recovery of the complete dataset. C) Sequence recovery of low shape complementarity interfaces. D) Sequence recovery of high shape complementarity interfaces. E) Sequence recovery of antigen-antibody complexes. Dashed lines represent median and triangles represent mean recovery values.

1. Standard Rosetta design (FixBB)
2. Surface-centric design (RosettaSurf)
3. Single-site-scanning surface-centric design (RosettaSurf-site)

All protocols operate on fixed backbones of the protein complexes that are presented in the native conformation that mediates the interaction. Consequently the sequence recovery success will largely depend on side-chain placement in a given backbone. The surface-centric design approaches are compared to the standard FixBB Rosetta design protocol, using the ref2015 scoring function (60). In the presented benchmarks, the Rosetta FixBB design protocol serves as baseline to assess the impact of surface-centric design on sequence recovery.

In a first step, the target protein’s interface is stripped off its native sequence by mutating all interface residues to alanine. Second, the different design protocols were employed on the interface positions with FixBB using the Rosetta energy function to select mutations while RosettaSurf was guided by our surface similarity scoring term (Fig. 2A). RosettaSurf was therefore provided with the native protein surface that we aimed to mimic. With this setup it is possible to evaluate the sensitivity of the Surf_S_ score to recover native amino acids compared to a known ground truth.

All different design protocols were employed for the bound and unbound states of the protein complexes and results are reported by their category as mean sequence recovery rate (Fig. 2A). Overall, sequence recovery rates were higher for surface-centric design pipelines while standard Rosetta sequence design showed lower recovery rates regardless of the presence of the binder (Fig. 2B). For all studied complexes, RosettaSurf and RosettaSurf-site outperformed FixBB by 36-39 percentage points in the unbound and 30-31 percentage points in the bound state. As expected, sequence recovery was in general more successful in the presence of the binding partner due to the reduced number of possible side-chains and rotameric conformations. However, even in the presence of the binder, surface-centric design can be applied to improve results. FixBB demonstrated improved performance for the antibody/antigen test set, however, the surface-centric protocols still reached better results that were comparable to the other categories. Worth noting is the 100% sequence recovery success of RosettaSurf-site for the D8 protein-vv138 antibody complex. The FixBB protocol performed slightly better for low-complementary (55%) than for high-complementarity complexes (48%), while RosettaSurf and RosettaSurf-site resulted in similar recovery rates of ∼85% in both cases with the notable exception of RosettaSurf-site for low-complementarity complexes. Here, RosettaSurf-site was able to recover 92% of the sequence and for one complex of that category, the Enterotoxin G – T-cell receptor complex, even 100%.

Furthermore, we analyzed more closely outlier decoys, i.e. structures that scored high in surface similarity but showed only little sequence recovery when designing with RosettaSurf. We set a threshold of up to 50% as low recovery and a Surf_S_ score of greater or equal to 0.9992 as this value marks the lower end of structures with high sequence recovery (>= 70%), leading to one candidate from the RSV epitope-scaffold-Motavizumab complex. When investigating which types of amino acids were common failures, we identified general geometric resemblance as likely reason. Several amino acids are frequently replaced by structurally similar residues, e.g. isoleucine by valine and lysine by arginine.

As for sequence recovery without the binder, RosettaSurf and RosettaSurf-site performed better for complexes of low shape complementarity with 81% and 80% recovery, respectively. However, sequence recovery for high-complementary and antibody/antigen complexes (66-76%) was still substantially higher than for FixBB (25% and 45%).

In addition, we investigated possible reasons for structures with low sequence recovery but high surface similarity scores. We adjusted the selecting criteria to structures with <= 30% sequence recovery and >= 0.9996 Surf_S_ score, observing similar results as for the bound benchmark. Four structures, all from the Colicin E9-IM9 complex fulfill these criteria. Again, mainly amino acids with similar geometrical features were common mismatches between recovered and native amino acids, e.g. phenylalanine, tyrosine, and histidine, aspartatic acid and asparagine, as well as valine and threonine.

Overall, similar performance of RosettaSurf and RosettaSurf-site showed that performing the combinatorial sampling of all possible rotamers simultaneously was not necessary. During the RosettaSurf-site pipeline amino acids were sampled individually at each design position and only a subset of the amino acids were used for combinatorial sampling based on the surface similarity score, resulting in similar recovery outcomes.

Our results have thus implications for the design of functional proteins as the high success of recovering natural protein interfaces may be promising for the design of proteins displaying defined surface properties, as is the case for immunogen design.

### Computational SSM screening to improve protein binding

A common problem in protein design is the optimization of PPIs to generate high affinity protein binders. Experimental screening methods, e.g. site-saturation mutagenesis (SSM) and combinatorial libraries, provide insights into mutations that improve binding interactions but are time and resource-consuming experiments. We tested if RosettaSurf-site could be a fast and efficient computational screening alternative to experimental based approaches. We studied the optimization of *de* novo designed interleukin-2/15 antagonists that bind specifically to the IL-2R*β γ*_*c*_ receptor (61). The computationally designed interleukins were optimized for binding to the interleukin receptor by performing SSM followed by combinatorial libraries based on the identified mutations. The described study did not only report sequence information but included additional structural characterization of the resulting design, thus allowing a fair comparison to our computational method. This level of data completeness is rare in many other design endeavors that have used this type of optimization strategy (39,62).

We performed RosettaSurf-site similar to the approach described above (see “Protein Interface Sequence Recovery”), however, with the selection criteria being shape complementarity of the design and IL-2R*βγ*_*c*_ interface rather than similarity as shape complementarity has been shown to be a key feature of high affinity binding interactions (63). RosettaSurf-site allows exhaustive computational sampling of amino acids similar to SSM experiments, screening all possible amino acid substitutions at the interface of the interleukin design. Our study is based on the reported crystal structure of the interleuking-2/15 design in complex with its cognate receptor (PDB: 6DG5). We selected interface positions that were tested during SSM and later included in the combinatorial library for our benchmark screening. Positions not interacting with the receptor, i.e. not within a distance of 5 Å, were excluded from the selection, resulting in a total of five positions. In our design protocol, the five selected residues were sampled with RosettaSurf-site and mutations were evaluated based on the shape complementarity to the receptor surface. The computational results were compared to the composition of the combinatorial library that was constructed based on the preceding SSM screening. We analyzed whether RosettaSurf-site was able to identify mutations that are present in the highest affinity binder as well as the performance of the Rosetta energy function to retrieve similar mutations. After performing all possible mutations with RosettaSurf-site, the four top-ranking amino acids for each position were selected (Fig. 3), in line with the maximum number of amino acids included in the combinatorial library.

**Figure 3.**
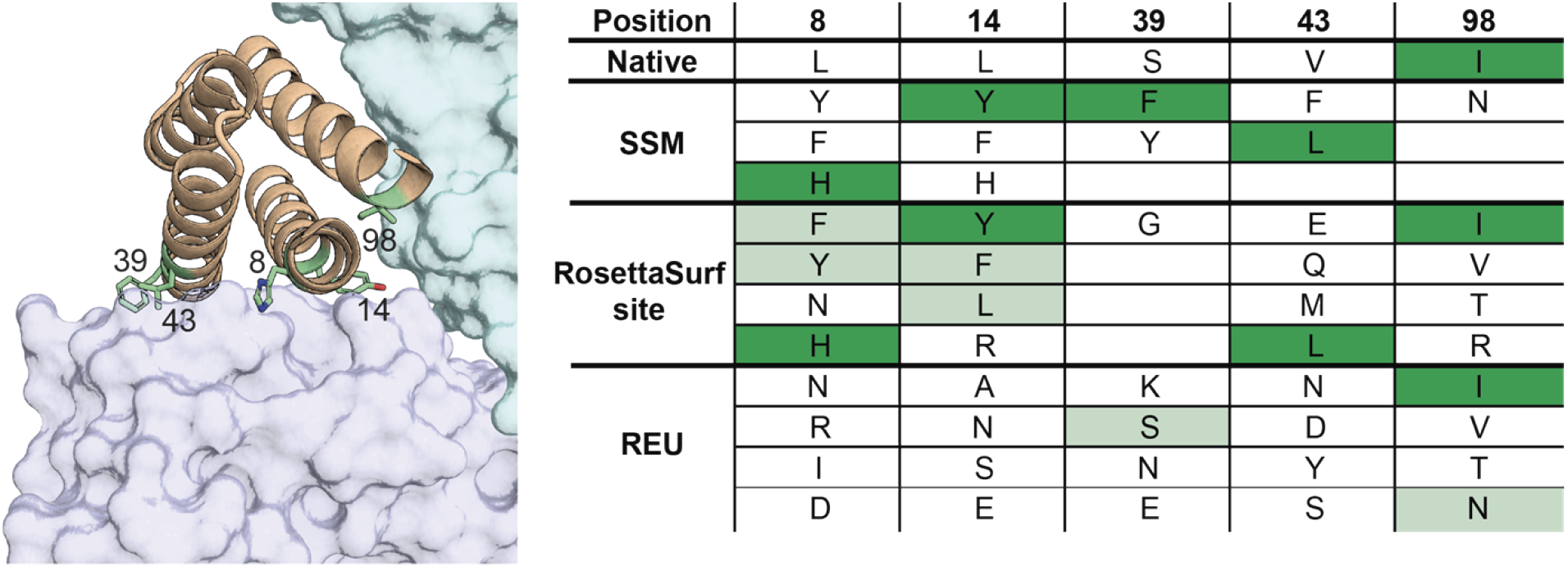
Comparison of SSM data obtained for the designed interleukin-2/15 antagonists in comparison to RosettaSurf-site predictions. The structure highlights the five selected positions of the interleukin design that were computationally and experimentally sampled. Different sampling results of the experimental SSM, RosettaSurf-site with SC, and Rosetta’s energy function are reported in the table. Mutations resulting in the experimentally reported best binding design are highlighted in dark green. RosettaSurf-site was able to recover four out of five key binding mutations (dark green) while evaluating mutations with Rosetta’s energy function could only retrieve one binding mutation at position 98. Additionally, RosettaSurf-site was able to recover four affinity improving mutations not present in the best binding design, whereas Rosetta’s energy function could identify only two of these mutations (light green).

RosettaSurf-site recovered four out of the five mutations present in the best design, with residue 39 being at the edge of the interface and contributing minimally to the protein-protein interaction, thus making it more challenging for this approach. Four additional residues included in the combinatorial library were recovered, that were observed to improve binding in the SSM but were not present in the highest affinity binder. In contrast, selecting mutations based on Rosetta energy alone recovers only a single mutation observed in the best binder and two residues present in the combinatorial library. These results show the potential of surface-based design of single point mutants as fast and promising approach to select candidates for combinatorial libraries to improve binding interactions without requiring preceding screening experiments like SSM.

### Surface-centric design of a novel RSV site 0 immunogen

In recent years, protein design has shown promising results in the field of immunoengineering, allowing the computational design of epitope-focused immunogens that were shown to elicit functional antibody responses in mice and non-human primates (34,36,37,42). To achieve an epitope-specific immune response, the epitope is transplanted from the viral antigen to an unrelated small protein scaffold, presenting the antigenic site in isolation. We applied the RosettaSurf protocol to the design of immunogens mimicking the antigenic site 0, an epitope on the F-glycoprotein of Respiratory Syncytial Virus (RSV) which consists of an irregular α-helix and a 10-residue long loop. Previous studies have shown that identifying scaffolds that can present such complex structural motifs is challenging, and specifically for site 0 they are currently not available in the known structure space. *De novo* design methods have been successfully used to build scaffolds from scratch (37,38), but the design process remains challenging and expertise in *de novo* protein design is critical. The RosettaSurf approach allows to rescue protein scaffolds that only present partial structural matches to the epitope structure motif.

As a demonstration of the use of RosettaSurf to transplant the epitope site into an unrelated scaffold, a two-step strategy was employed: 1) side chain grafting of the α-helical segment onto a small, monomeric protein scaffold (Fig. 4A); 2) RosettaSurf design of the remaining antigenic site, including the surface generated by the epitope loop and transition to the helical fragment (Fig. 4A). The grafted helical segment serves as anchor point around which the surface can be optimized to mimic the complete antigenic site.

**Figure 4.**
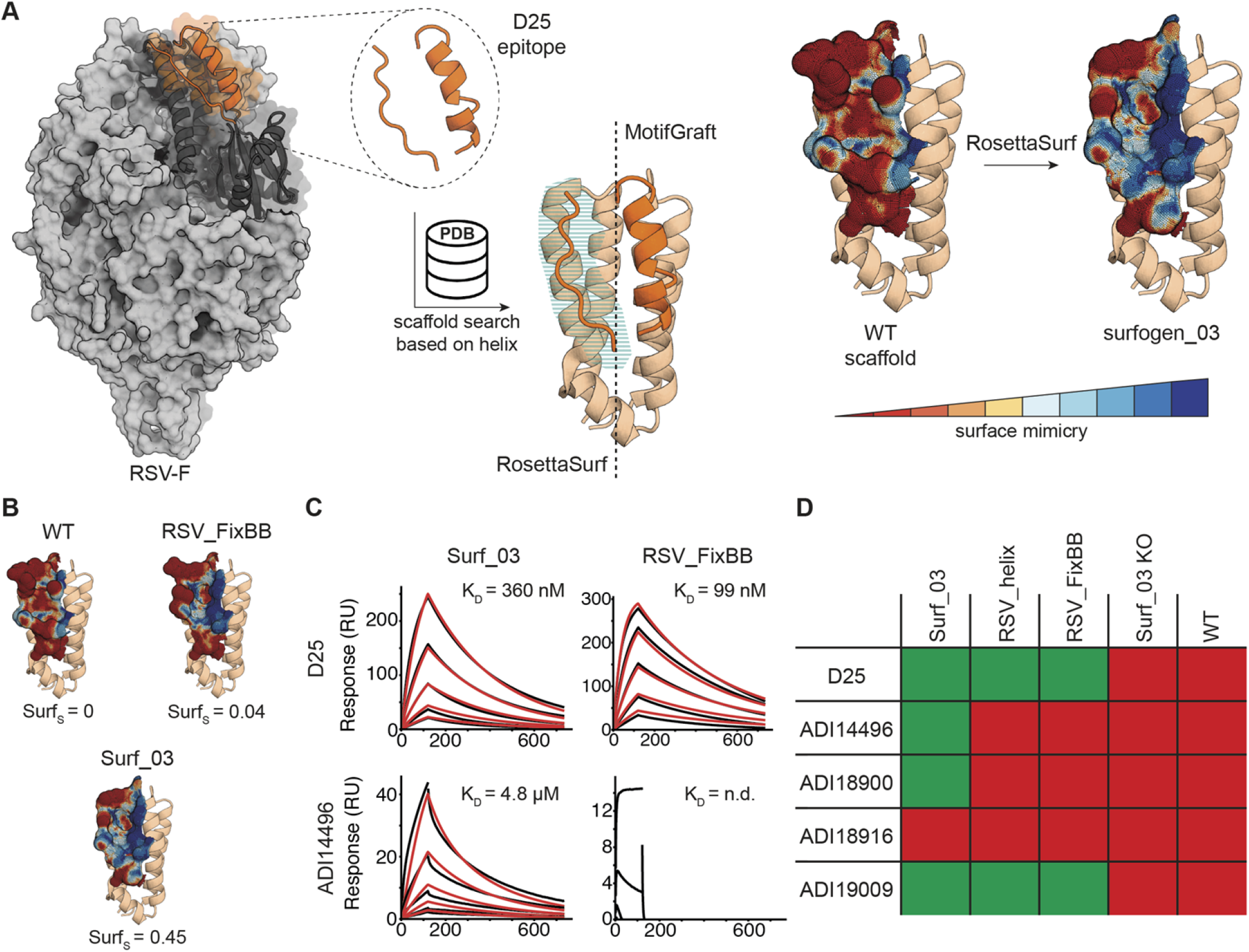
Surface centric design of a viral antigenic site present the in RSVF. A) Design process of site 0-mimicking protein scaffolds. Starting scaffolds are selected from the PDB based on structural alignments with the epitope helix. The surface mimicking designs are generated by grafting the side chains of the helix segment of the epitope onto the scaffold and surface-centric design is employed to optimize the loop region. Before and after design of the surface compared to native site 0. Blue areas indicate high similarity. B) Mimicry of surface geometry of WT scaffold, RSV_FixBB, and Surf_03 designs compared to native site 0. C) Representative SPR measurements of Surf_03 and RSV_FixBB against site 0-specific antibodies D25 and ADI14496. D) Binding profiles of Surf_03, a FixBB designed protein, and a helix-only design against a panel of site-specific antibodies with green indicating binding and red cells corresponding to non-binding. A knockout mutant of Surf_03 and the WT protein binding profiles are listed as reference.

**Figure 6.**
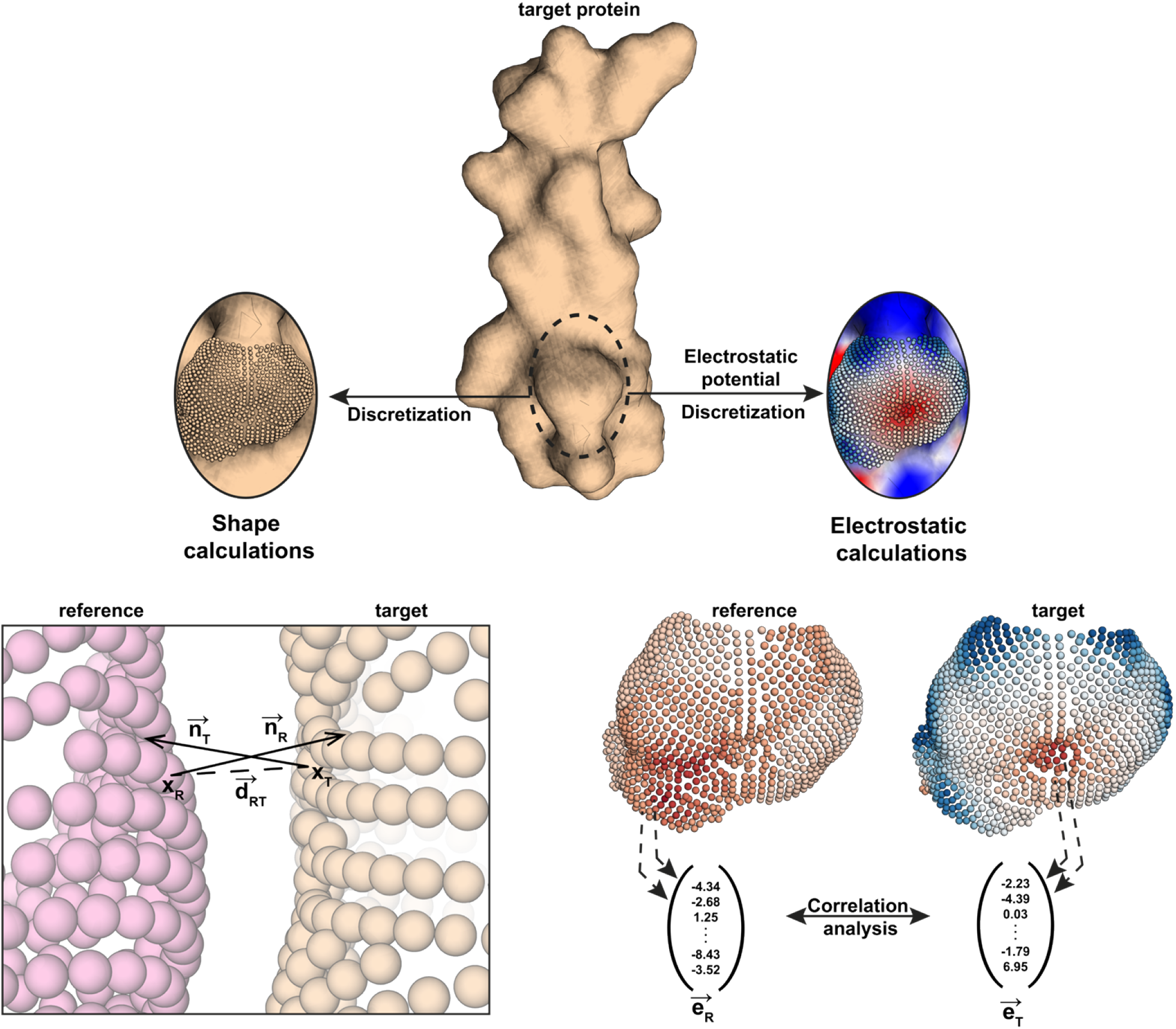
Computation of the surface similarity score (Surf_S_). Protein surface is converted into a point cloud where each point is used to compute shape and electrostatic features. To compute the Surf_S_ score, all individual points of the point clouds are compared and the shape similarity value is derived from closest points of the two surfaces in space while electrostatic similarity is evaluated by correlation analysis of the electrostatic potentials of both surfaces.

We identified the NarX histidine kinase receptor (PDB: 3EZI) (59) as promising scaffold. The small, monomeric protein can accommodate the epitope α-helix (Cα RMSD: 0.4 Å) and offers a sufficiently large surface area around the helix to be optimized with RosettaSurf. We generated 760 designs using the RosettaSurf pipeline and the site 0 surface, as observed in complex with site-specific antibody D25 (57), as a sequence optimization target. The best design was selected based on the highest surface similarity score and was further optimized with six mutations (Surf_03). First, five point mutations, which were not part of the interface, were introduced to resolve clashes between native scaffold and epitope residues in the α-helix as well as steric hindrance with residues introduced during surface-centric design. In addition, we mutated a polar residue in the binding pocket to a hydrophobic residue based on the sequence profile of the top 20 surface-designed decoys.

Two additional variants of the same protein scaffold were designed to test whether there were advantages in using RosettaSurf. One design containing only the α-helix of site 0 (RSV_helix) and three point mutations that resolve clashes with the side chain grafted epitope helix, similar to Surf_03. The second design was generated with FixBB (RSV_FixBB), starting with side-chain grafting of the epitope α-helix, and allowing to design the same amino acid positions as those of Surf_03. The RosettaSurf design (Surf_03) reaches a surface similarity score of 0.45 compared to the antigenic site in the native viral protein RSVF, while the native scaffold scores 0. The helix-grafted base-design scores 0.06 and the RSV_FixBB design 0.04 in surface resemblance.

To test experimentally our predictions, the designs were expressed and purified, and the binding affinities were measured using a panel of monoclonal antibodies that engage the site 0 epitope (Fig. 4B) (37,57,64). In essence, this panel of monoclonal antibodies (mAbs) was used as conformational probes to assess the surface mimicry presented by the designs. We compared the binding profiles of the three designs, a negative control (Surf_03 KO) and the WT scaffold using surface plasmon resonance (Fig. S3). All three designs bound to the D25 and ADI19009 mAbs, indicating that these mAbs mostly rely on the helical segment of the epitope. Surf_03 was the only design recognized by two additional antibodies (ADI14496 and ADI18900), indicating that the higher surface mimicry achieved through RosettaSurf design improves the presentation of the antigenic site and promotes binding of additional site-specific antibodies. The designed immunogen demonstrates RosettSurf’s capabilities to sculpt protein surfaces with a high degree of accuracy which could be of use to introduce functional sites into protein scaffolds by optimizing the molecular surface.

## Discussion

In this work we propose a surface-centric protein design approach and demonstrate its accuracy in several *in silico* benchmarking and experimental design tasks. The protocol can either be used to optimize surface similarity or surface complementarity depending on the design task at hand. Our ideas to leverage the surface geometrical and chemical features stem from observations that surfaces displaying similar patterns can have similar functional roles (e.g. in PPIs (2,11,14,65)). Technically, our framework is derived from earlier work by Lawrence and Coleman that introduced a fast and efficient way to evaluate shape complementarity as well as work by McCoy et al. that addressed electrostatic complementarity in interfaces. Based on some of these principles, we further implemented a surface similarity score to evaluate both shape and chemistry of surfaces. The resulting surface similarity score is implemented with a focus on surface-centric protein design pipeline, allowing easy interaction and modification of the algorithm inside Rosetta design protocols.

We showed the high accuracy of the surface score in recovering single amino acid identities by their surface properties and show its benefit when used in conjunction with the Rosetta energy score. Further, we highlight the use of surface similarity score inside a Monte Carlo sampling approach and its performance in the sequence recovery of complete protein interfaces. The incorporation of surface similarity clearly increases sequence recovery rates relative to other scoring schemes.

This approach can be readily applied for the design of experimental combinatorial libraries to reduce library sizes, as the high sequence recovery rates suggest that surface-centric design is especially interesting to design novel proteins that mimic surface patches of other proteins, as it is the case for the design of novel immunogens for vaccine development (36–38).

We demonstrated that surface-centric design could be used to generate targeted libraries to optimize binding specificity of a previously reported IL-2 design (61). With the ranking of all mutations, we were able to recover amino acid variants that were also obtained by experimental screening through a saturation mutagenesis library. This approach represents a straightforward computational screening method to detect mutations that modulate specificity and affinity in protein-protein interactions. Specifically, this strategy is a fast and accessible alternative to experimental screening techniques.

Finally, we applied the surface-centric design protocol to engineer novel immunogens to present the surface patch of an antigenic site present in the RSVF protein. We used a structurally complex epitope (site 0 from RSVF), that consists of two structural segments, as an example for structurally challenging sites that remain difficult for computational design approaches. The surface-centric designs show a broader binding profile across a panel of monoclonal antibodies as compared to other design approaches, suggesting that the surface presented is a closer mimic of the native antigenic site.

Possible applications of surface-centric design range from the computational design of highly specific protein-protein interactions, focusing on optimizing the surface complementarity both in shape and chemistry. On the other hand, surface similarity enables the computational design of proteins that may recapitulate precise surface features of target surfaces, which could then find applications in immunogen design, where surface similarity allows the design of immunogens mimicking an antigenic site of interest that could ultimately be used as probes for antibody isolation or vaccine antigens.

Ultimately, we introduce a new conceptual approach in computational protein design where the fine features of molecular surfaces are explicitly optimized. Through this route it will now be possible to explore the tantalizing hypothesis of the existence of a “surface degeneracy code”, which in some cases may allow to represent similar surface patches using very distinct sets of amino acids. Importantly, this capability could also enable the presentation of similar surface patches in proteins with completely different backbone architectures which could again realize many endeavors in functional protein design that are thus far out of reach.

## Materials and Methods

### RosettaSurf framework

The RosettaSurf protocol operates at the solvent-excluded surface level of a protein structure, and its core operation is the comparison between surfaces. To compare molecular surfaces, we defined a score that quantifies the similarity between two surfaces considering both shape and electrostatic features, and incorporated it within the Rosetta software package. The molecular surface is generated from the three-dimensional atomic coordinates of a protein (4,5) and stored as a discrete point cloud (Fig. 5). Representation of the surface as a point cloud allows the featurization of the points with geometrical and chemical descriptors (Fig. 5) and enables rapid calculations of the surface similarity score. We refer to the mutable surface as target while the reference surface used for comparisons is denoted as reference.

To describe the surface geometry, we developed a descriptor based on concepts introduced by Lawrence and Colman (19) that quantifies shape relationships of two surfaces relying on normal vector comparisons that are derived at each point of the surface. To evaluate surface shape similarity, two protein structures are first aligned and the closest surface points between target and reference are identified. Next, the normal vectors of the surface points are compared by first computing their dot product to obtain the enclosed angle, followed by distance-based scaling that penalizes points that are far apart (Fig. 5). Each comparison yields a shape similarity score for the considered pair of points and the similarity of the entire surface results from computing the mean of all pair-wise point comparisons. Since the identification of closest points depends on the starting surface, the surface scores are not identical depending on which structure is the considered the reference and the target and thus the scores are computed in both combinations. A robust surface similarity score is obtained by considering the similarity of the target surface compared to the reference surface and vice versa, effectively averaging the shape similarity to correct for differences during the selection of closest points.

Electrostatic similarity (E_S_) of two surfaces is derived from comparing their electrostatic potentials, i.e. E_S_ is assessed by computing the correlations of two electrostatic fields (Fig. 5). To quantify surface electrostatics, the electrostatic potentials are computed over the continuous electrostatic field and discrete charge values are assigned to every point of the surface using the APBS software (46) (Fig. 5). The representation of the target and reference surface point clouds as vectors allows fast computation of Pearson’s and Spearman’s rank correlations as described by McCoy and colleagues (20). The resulting correlation coefficients capture the similarity of the individual potential values as well as the overall trends in electrostatic similarity of the two surfaces (18,20).

To accurately capture the relative contributions of both shape and electrostatics on the similarity of molecular surfaces, we combined both scores into a single surface similarity score (Surf_S_). Since we were interested in assessing similarity of surface sites that are well maintained and relevant to functional protein design, we focused on interfaces of PPIs and performed logistic regression on a dataset of 2,660 protein complexes to optimize the individual weights of the Surf_S_ score for each component, shape and electrostatics. For each amino acid type (excluding cysteines) a total of 140 complexes containing the respective amino acid in the interface region were considered. All possible point mutations to non-native amino acids were generated and their geometric and electrostatic similarity to the native surface measured, resulting in a dataset containing shape and electrostatic properties for each mutation. Logistic regression was applied to identify the optimal set of weights to combine shape and electrostatic features to optimize for the highest recovery rates of the native amino acids.

The Surf_S_ score is defined as follows:

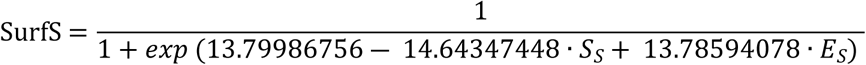

The final Surf_S_ score combines both properties into a single score scaled from 0 to 1, where 1 represents highly similar surfaces. We note that the developed surface similarity score can be straightforwardly converted to surface complementarity (S_C_) by inverting the normal vectors of the reference surface. Complementarity has been typically used to evaluate PPIs and was first demonstrated by Lawrence and Coleman for molecular shapes (19).

### Surface-centric protein design protocol

We developed the RosettaSurf design protocol, as part of the RosettaScript software environment (47), which utilizes the described surface scoring function during the sequence sampling stage of the design process to bias the selection of amino acids and rotamers towards a desired surface configuration in terms of geometric and electrostatic properties (Fig. 2A). Thereby, it is possible to design proteins for function without relying on protein grafting but by optimizing the molecular surface, for instance, to mimic a given active site.

With the RosettaSurf design protocol we explicitly optimize molecular surface features during the protein design process. In practice, RosettaSurf performs sequence design in a way where the mutable surface of the design scaffold (target) is optimized to closely match the surface features of a reference protein (reference) (Fig. S1). To efficiently explore the sequence space during the design process, Monte Carlo simulated annealing guides the optimization of rotamers, where substitutions of residues are scored based on the resulting surface and accepted if they pass the Monte Carlo criterion that is implemented as the Surf_S_ score.

To reduce computational time spent on rotamer sampling in a combinatorial fashion on the overall surface, we implemented a single amino acid scanning surface-centric protein design approach (RosettaSurf-site). This protocol samples amino acids individually at each position of the design surface, selects the top three rotamers according to the surface and samples those combinatorially with other designable positions, reducing the combinatorial possibilities.

### Benchmark datasets

The dataset of the single amino acid recovery benchmark consists of 1,900 protein complexes, with 100 complexes for each amino acid identity except cysteine. We divided each of these protein complexes into target proteins, where the interface can be mutated, and binders, the proteins that serve as context for the target. The interface of the target protein is defined as residues that are within 7 Å *C*_*β*_-distance of the binder and have the *C*_*β*_-atom pointing towards the binder. This selection ensures that the amino acid side chains are part of the interface and the contribution of the residue to the binding interaction is not solely due to backbone interactions. The full target interface is converted to alanine, effectively removing any side chain memory of the native structure that would restrict the placement of new rotamers and introduce biases towards the native sequence. Cysteine residues are ignored as they can form chemical linkages in the form of disulfide bonds, thus generating a surface that is not attributable to an individual residue.

To test the performance of the surface scoring function we sought to perform a benchmark to evaluate sequence recovery at the interfaces of protein-protein interactions with varying shape complementarity. We assembled a diverse dataset consisting of nine protein complexes capturing different aspects of PPIs and grouped them into three different categories: 1) low-shape complementary interactions which include Enterotoxin G – T-cell receptor complex (PDB: 3MC0) (48), Ribonuclease A in complex with its inhibitor (PDB: 1DFJ) (49) and domain 2 of VEGFR1 in complex with PIGF (PDB: 1RV6) (50); 2) High-shape complementarity interactions which include the complex between Colicin E9 and IM9 (PDB: 1EMV) (51), Bovine beta-trypsin in complex with CMTI-I (PDB: 1PPE) (52) and PD-L1 in complex with a nanobody (PDB: 5JDS) (53); 3) antigen-antibody interactions which include HIV-gp120 in complex with CD4-binding site antibody b13 (PDB: 3IDX) (54), RSV epitope-scaffold in complex with Motavizumab (PDB: 4JLR) (34) and the Vaccinia virus D8 protein in complex with the antibody vv138 (PDB: 6B9J) (55). The shape complementarity of the complexes in the first two categories was assessed by the Rosetta shape-complementarity filter. Protein complexes with a shape complementarity score less than 0.65 were classified as low-complementarity. During the benchmark analysis we distinguish between bound and unbound proteins, where unbound proteins are obtained by removing the protein binder from the holo-crystal structure.

Data analysis was performed with the help of the rstoolbox Python library (56).

### Computational design of proteins mimicking the antigenic site 0 in RSV

The structure of antigenic site 0 was extracted from the crystal structure of prefusion stabilized RSVF in complex with antibody D25 (PDB: 4JHW) (57). The D25 epitope consists of an irregular α-helix (residues 196-209) and a 10-residue loop (residues 61-70). To identify putative scaffolds, we performed a structural search based on the irregular α-helix against 55,574 monomeric, helix-containing crystal structures from the Protein Data Bank (PDB, from September 2015) using Rosetta’s MotifGraft algorithm (58). Matches were filtered at a backbone RMSD threshold below 0.55 Å and less than ten atomic clashes at the interface, resulting in 13 scaffold candidates. After visual inspection, we selected the NarX histidine kinase receptor (PDB: 3EZI) (59) as scaffold, a 107-residue long, monomeric protein that aligned to the epitope helix with a C_α_ RMSD of 0.4 Å and provided additional surface area to mimic the entire antigenic site.

In a first step, we transplanted the side chains of the epitope helix contacting the D25 antibody onto the selected scaffold using MotifGraft, followed by the introduction of three mutations on the protein scaffold that were not part of the interface to resolve steric clashes with the transferred side chains, resulting in the design RSV_helix. Subsequently, we performed surface-centric design on 19 residues using the RosettaSurf pipeline to increase surface mimicry of the site 0 around the epitope helix, resulting in 760 design decoys. Surface-centric design was performed in the presence of the D25 antibody and site 0 of RSVF served as a reference to which our designed surface was optimized. We selected the decoy with the highest surface mimicry score, named surf_01, and optimized the design in additional steps. To allow accurate display of the designed surface, we introduced two point mutants in the scaffold adjacent to the optimized surface patch to avoid steric hindrance (surf_02). Lastly, after comparing the similarity of surf_02 and the native antigenic site, we identified a lysine residue at position 18 in our design with suboptimal mimicry. Evaluating the sequence profile of the 20 best scoring decoys sorted by surface mimicry revealed a strong preference for leucine at this position and the residue was incorporated into a new design (Surf_03). Finally, we designed a version of RSV_helix that used Rosetta’s fixed-backbone design (FixBB) to serve as comparison to our surface-optimized designs. We designed 1’000 decoys in the presence of D25 antibody, allowing mutations to occur at the same 19 residues as was the case for RosettaSurf design. All designs converged to an identical sequence which was selected as design RSV_FixBB. Based on Surf_03 we designed a knockout mutant (Surf_03_KO) with a N74Y mutation in the epitope helix.

### Protein expression and purification

#### Designs

Genes for all designs were purchased as DNA fragments from Twist Bioscience, and cloned into pET11 vectors, containing a N-terminal MBP-tag and His-tag as well as a TEV cleavage site, for bacterial expression. Plasmids were transformed into *E. coli* BL21 (DE3 pLysS) (Merck, #69451-3) and grown overnight in LB media at 37 °C. Pre-cultures were diluted 1:50 and inoculated to an OD_600_ of 0.6 in terrific broth (Condalab, #PRO1246.05) at 37 °C and expression was induced by the addition of 1 mM isopropyl-β-D-thiogalactoside (IPTG). Cultures were harvested after 18-20 hours at 20 °C. Pellets were resuspended in lysis buffer (50 mM Tris, pH 7.5, 500 mM NaCl, 5% Glycerol, 1 mg/ml lysozyme, 1 mM PMSF, and 1 μg/ml DNase) and sonicated on ice for a total of 12 minutes, in intervals of 15 s sonication followed by 45 s pause. The lysates were cleared by centrifugation (48’384 g, 20 min) and purified using a His-Trap FF column on an Äkta pure system (GE Healthcare), followed by size exclusion on a HiLoad 16/600 Superdex 75 column (GE Healthcare) in phosphate-buffered saline (PBS). Protein concentrations were determined by measuring the absorbance at 280 nm on a Nanodrop (Thermo Scientific). The designed proteins were concentrated by centrifugation (Millipore, #UFC900324) to 1 mg/ml, snap frozen in liquid nitrogen, and stored at −80°C.

#### Antibody variable fragments (Fabs)

For Fab expression, heavy and light chain DNA sequences were purchased from Twist Biosciences and cloned separately into the pHLSec mammalian expression vector (Addgene, #99845) using AgeI and XhoI restriction sites. Expression plasmids were premixed in a 1:1 stoichiometric ratio, co-transfected into HEK293-F cells, and cultured in FreeStyle medium (Gibco, #12338018). Supernatants were harvested after 1 week by centrifugation and purified using a kappa-select column (GE Healthcare). Elution of bound proteins was conducted using 0.1 M glycine buffer (pH 2.7), and eluates were immediately neutralized by the addition of 1 M Tris ethylamine (pH 9), followed by buffer exchange to PBS (pH 7.4).

#### Binding affinity determination by surface plasmon resonance (SPR)

SPR measurements were performed on a Biacore 8K (GE Healthcare) with HBS-EP+ as running buffer (10 mM HEPES pH 7.4, 150 mM NaCl, 3 mM EDTA, 0.005% v/v Surfactant P20, GE Healthcare) at room temperature. Approximately 700 response units (RU) of Fabs were immobilized on a CM5 sensor chip (GE Healthcare) via amine coupling, and designed monomeric proteins were injected as analyte in two-fold serial dilutions. The flow rate was 30 μl/min with 120 s of contact time followed by 400 s dissociation time. After each injection, surface was regenerated using 0.1 M glycine at pH 3.5. Data were fitted using 1:1 Langmuir binding model within the Biacore 8K analysis software (GE Healthcare #29310604).

## Author contributions

A.S. and B.E.C. conceived the work and designed the experiments. A.S. implemented the described protocol with input from J.B. and performed computational design simulations and benchmark analysis. S.R. expressed and purified the reported proteins and performed SPR assays. A.S., M.D., and A.L. developed the surface similarity score. A.S. and B.E.C wrote the paper. All authors commented on the manuscript.

## Acknowledgments

We would like to thank Pablo Gainza, Fabian Sesterhenn, Che Yang, and Sarah Wehrle for helpful discussions and support regarding *in vitro* experiments. We thank Sailan Shui, Freyr Sverrisson, and Arne Schneuing for comments on the manuscript. We also thank Luki Goldschmidt and Brian Coventry for implementing the shape and electrostatic complementarity evaluation within Rosetta, respectively. Additionally, we would like to acknowledge the high-performance computing facility (SCITAS) for their technical support. We would also like to acknowledge the Swiss National Supercomputing Centre (CSCS) for their support in computing time. This work was supported by the European Research Council (starting grant no. 716058), the Swiss National Science Foundation (grant no. 310030_188744), the NCCR Molecular Systems Engineering and the NCCR Chemical Biology. Andreas Loukas was supported by the Swiss National Science Foundation project “Deep Learning for Graph-Structured Data” (grant number PZ00P2 179981). Jaume Bonet was funded by the EPFL Fellows postdoctoral fellowship. The funders had no role in study design, data collection and analysis, decision to publish, or preparation of the manuscript.

## Data availability statement

All data and scripts necessary to recreate the analysis and design simulations described in this work are available at https://github.com/LPDI-EPFL/RosettaSurf.

